# Evaluation of the RNA-dependence of PRC2 binding to chromatin in human pluripotent stem cells

**DOI:** 10.1101/2023.08.17.553776

**Authors:** Yicheng Long, Taeyoung Hwang, Anne R. Gooding, Karen J. Goodrich, Skylar D. Hanson, Tenaya K. Vallery, John L. Rinn, Thomas R. Cech

## Abstract

Polycomb Repressive Complex 2 (PRC2), an important histone modifier and epigenetic repressor, has been known to interact with RNA for almost two decades. In our previous publication (Long, Hwang et al. 2020), we presented data supporting the functional importance of RNA interaction in maintaining PRC2 occupancy on chromatin, using comprehensive approaches including an RNA-binding mutant of PRC2 and an rChIP-seq assay. Recently, concerns have been expressed regarding whether the RNA-binding mutant has impaired histone methyltransferase activity and whether the rChIP-seq assay can potentially generate artifacts. Here we provide new data that support a number of our original findings. First, we found the RNA-binding mutant to be fully capable of maintaining H3K27me3 levels in human induced pluripotent stem cells. The mutant had reduced methyltransferase activity in vitro, but only on some substrates at early time points. Second, we found that our rChIP-seq method gave consistent data across antibodies and cell lines. Third, we further optimized rChIP-seq by using lower concentrations of RNase A and incorporating a catalytically inactive mutant RNase A as a control, as well as using an alternative RNase (RNase T1). The EZH2 rChIP-seq results using the optimized protocols supported our original finding that RNA interaction contributes to the chromatin occupancy of PRC2.

---

Many chromatin-associated proteins, including the epigenetic silencing complex Polycomb Repressive Complex 2 (PRC2), also bind RNA. The contributions of RNA in inhibition and/or recruitment of PRC2 are subjects of ongoing research. In our recent study (Long, Hwang et al. 2020), we concluded that RNA was required for PRC2 occupancy on chromatin in human induced pluripotent stem cells. Recently, researchers have contacted us, questioning (1) if our RNA-binding mutant of PRC2 is in fact a separation-of-function mutant, or instead has lost histone methyltransferase (HMTase) activity, and (2) whether our rChIP-seq (RNase A-Chromatin Immunoprecipitation followed by deep sequencing) method is compromised by RNase-induced precipitation of chromatin leading to a global gain of non-targeted DNA (Healy, Zhang et al. 2023). After our publication, we continued to study our experimental reagents and methods. We now update the community on these new results so they can evaluate the extent to which our previous conclusions might require modification and how the rChIP-seq method can be improved.

## RNA-binding mutant of PRC2

Our RNA-binding mutant altered ten amino acids on the surface of the EZH2 subunit of PRC2, in two regions that had previously been implicated in RNA binding. We replaced the endogenous *EZH2* genes with this mutant by CRISPR genome editing of human iPSCs and selected two clones for further analysis. For comparison, we genome-edited the wild-type *EZH2* sequence into the endogenous *EZH2* genes to control for effects of the CRISPR genome editing. Using western blot analysis, we found that the levels of EZH2 and other PRC2 subunits were comparable in the mutant and wild-type iPSCs, and the H3K27me3 mark deposited by PRC2 was also maintained at comparable levels (Fig. 2b of (Long, Hwang et al. 2020)). ChIP-seq analysis with an anti-EZH2 antibody identified 247 genes with significantly decreased PRC2 occupancy, but most genes maintained their normal low PRC2 occupancy (Fig. 2f of (Long, Hwang et al. 2020)). Thus, in our CRISPR genome-edited iPSC lines, our data do not support the mutant EZH2 as having a global loss of function in expression, assembly, or HMTase activity.

If our RNA-binding mutant were in fact a loss-of-function mutant, it should have properties resembling those of true loss-of-function mutants. We therefore constructed true loss-of-function PRC2 iPSC lines for comparison. To this end, four poly(A) sites (three copies of SV40 poly(A) and one copy of bGH poly(A) sites) were inserted immediately downstream of the AUG translational start site of the endogenous *EZH2* genes (beginning of exon 2) or *SUZ12* genes (middle of exon 1) in our iPSC line. Several independent clones of cells were analyzed, and they all showed little or no residual H3K27me3 signal (Fig. 4 of (Long and Cech 2021)). In addition, these cells only continued growing for five passages and the residual surviving cells spontaneously differentiated, unlike our RNA-binding mutant EZH2 cells. The phenotypes of the EZH2-knockout and SUZ12-knockout cell lines were therefore distinct from those of our RNA-binding mutant EZH2 cell lines, supporting the conclusion that our mutant was not substantially compromised in HMTase activity in living cells. Furthermore, these results argue against the possibility that EZH1, which was not knocked-out in our EZH2 mutant cell lines, might have compensated for hypothetical EZH2 loss of function. If there is any compensation by EZH1 in the EZH2- or SUZ12-knockout cell lines, it is insufficient to restore substantial levels of H3K27me3.

We also re-tested our RNA-binding mutant PRC2 for HMTase activity in vitro. Designing such in vitro experiments presents a challenge, in that one must choose a form of PRC2 (PRC2.1 or PRC2.2, inclusion of accessory subunits, choice of splice isoforms) and also choose a form of the histone 3 substrate (size of the nucleosome array, pre-existing modifications or unmodified). In making these choices, one realizes that no decision will recapitulate the situation in vivo, where multiple forms of PRC2 are acting on multiple forms of chromatin. Although our published in vitro data were mostly performed with a PRC2 4-mer complex containing the core subunits EZH2, SUZ12, EED and RBPP4, we have now focused on a PRC2.2 5-mer complex that includes the short splice isoform of AEBP2. We chose this 5-mer complex, which assembles well when expressed in insect cells, because it has greater HMTase activity than the 4-mer complex (providing more accurate quantification) and is plausibly more biologically relevant.

HMTase assays with unmodified H3 protein and with commercial human cell polynucleosomes, which contain a mixture of natural modifications, both showed that the RNA-binding mutant had activity comparable to that of wild-type PRC2 (Fig. 1A). With reconstituted unmodified trinucleosomes as a substrate, on the other hand, the mutant produced less H3 methylation at shorter incubation times but then caught up upon prolonged incubation (Fig. 1B). This kinetic effect occurred with trinucleosomes reconstituted either by the dilution method (Long, Hwang et al. 2020) or by dialysis (Fig. 1C). Quantification of five independent experiments is shown in Fig. 1D.

**Figure 1.**
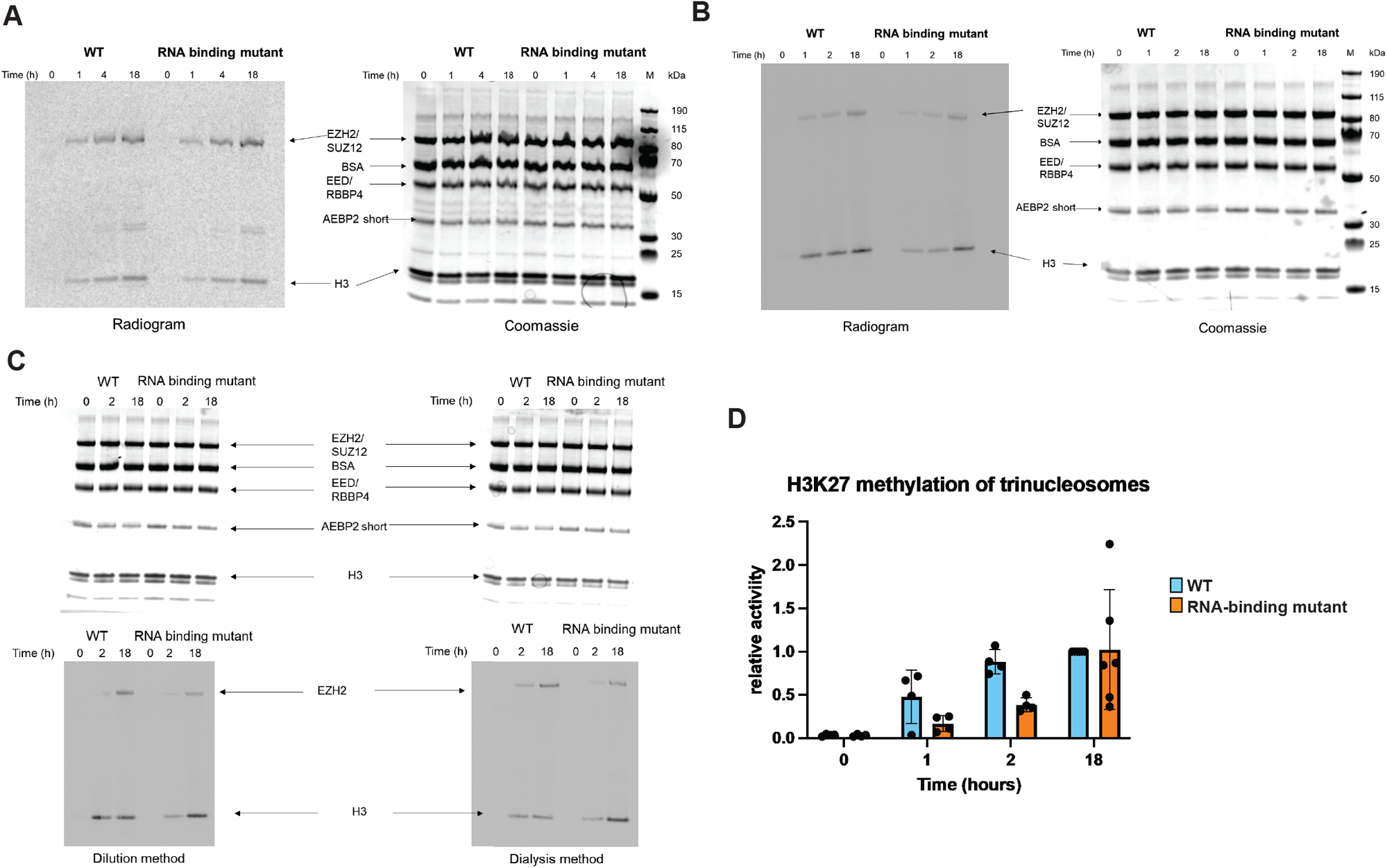
Activity assays show EZH2 RNA-binding mutant retains HMTase activity but differs from wild-type EZH2. **(A)** 5-mer PRC2 complex incubated with polynucleosomes and ^14^C-S-adenosylmethionine for the indicated times, then analyzed by SDS-PAGE. Radiogram shows EZH2 automethylation and H3 methylation. Coomassie staining of same gel shows equivalent loading of PRC2 and nucleosomal histones. **(B)** 5-mer PRC2 incubated with trinucleosomes reconstituted with 621 base-pair DNA and human histone octamer. **(C)** HMTase assays comparing trinucleosomes reconstituted by dilution or by dialysis. **(D)** Quantification of biological replicates of HMTase assays (5-mer PRC2, reconstituted trinucleosome substrate) performed over a period of three months. Data are normalized to the amount of H3 methylation obtained for WT PRC2 after 18 hours. Bars represent averages of the individual experimental values (dots).

**Figure 2.**
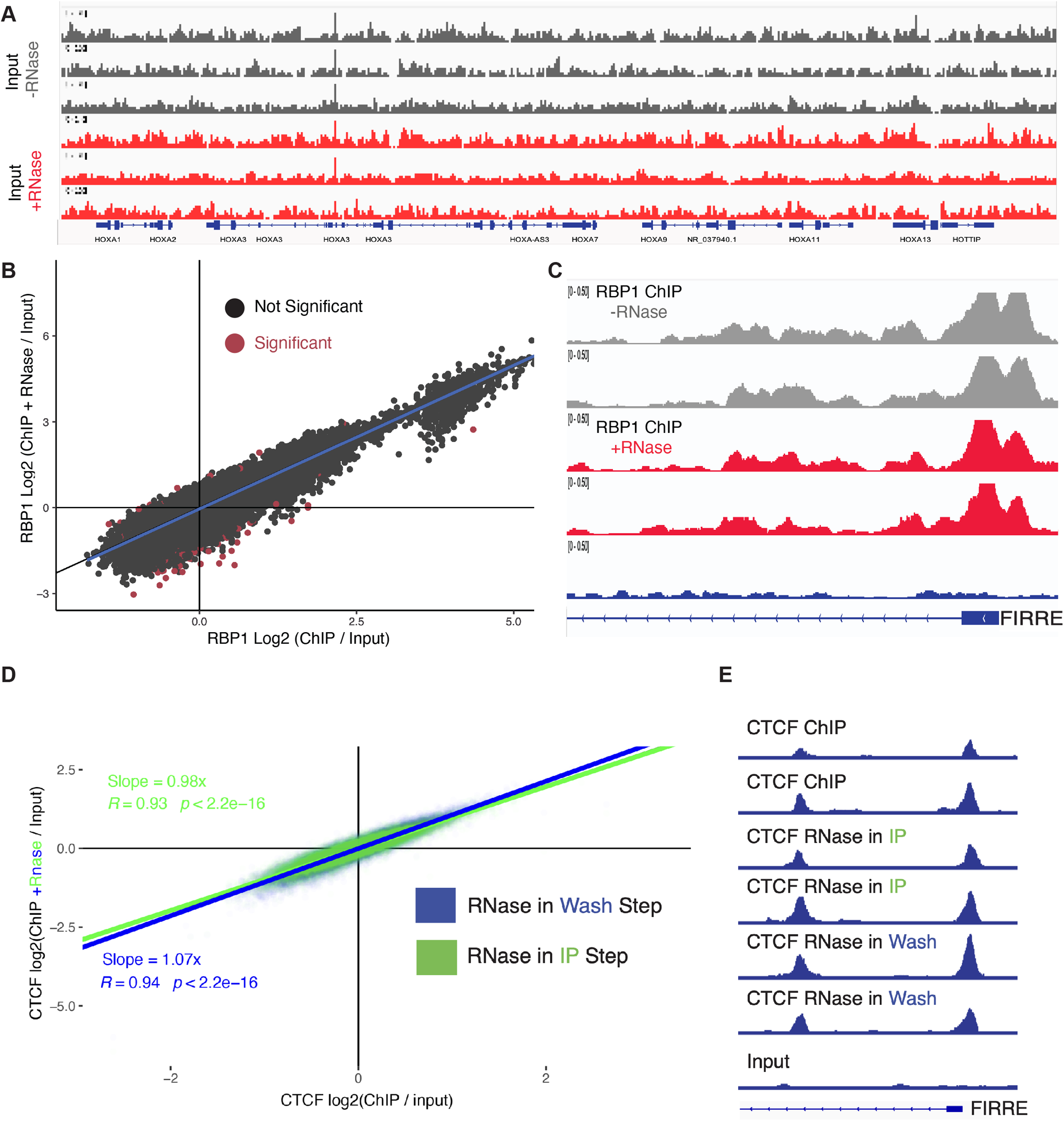
Testing rChIP-seq for differential DNA recovery and with a different cell line and different antibodies. **(A)** Genome tracks of sequences represented in input DNA with or without RNase A treatment according to Long, Hwang et al. 2020. Three replicates of each are shown. **(B)** rChIP vs. ChIP for a control protein, RBP1, which does not display RNA-dependent binding to chromatin, in K562 cells. Significant: MACS3 qVal < 0.05. **(C)** Example of RBP1 ChIP vs. rChIP over the FIRRE lncRNA locus. **(D)** rChIP vs. ChIP for CTCF in K562 cells, and data with a variation of the rChIP protocol in which the RNase A is added in the wash step instead of the IP step. **(E)** Example of CTCF peaks over the FIRRE locus obtained with different ChIP and rChIP protocols.

In conclusion, we found our RNA-binding mutant to be fully capable of maintaining H3K27me3 levels in human pluripotent stem cells. In our biochemical assays with the PRC2 5-mer complex, the RNA-binding mutant was fully active on polynucleosomes and histone H3 substrates, although slower methylation kinetics were observed with a trinucleosome substrate.

### The rChIP-seq method for evaluating RNA-dependence of chromatin binding

Testing the RNA dependence of a protein’s association with chromatin using RNase treatment is not a new idea. ChIP experiments with and without RNase treatment were used by the Rosbash laboratory in 2004 and have been employed by multiple groups since then (Abruzzi, Lacadie and Rosbash. 2004, Bernstein, Duncan et al. 2006, Jeon and Lee 2011, Thompson, Dulberg et al. 2015, Casale, Cappucci et al. 2019, Skalska, Begley et al. 2021). The new feature of rChIP-seq was to compare RNase-treated and untreated immunoprecipitated DNA genome-wide rather than focusing on single sites (Long, Hwang et al. 2020). Because RNase treatment is such a common tool, it is important that the technique be robust and reproducible not just for our PRC2 experiments, but for the field more generally.

Since the publication of rChIP-seq, we have performed additional experiments to test whether RNase A influences the properties of the protocol. These include testing whether RNase A perturbs the input chromatin to which all the ChIP signals are compared, whether rChIP-seq gives concordant results among cell lines, and whether other protein-antibody combinations are influenced by RNase A treatment. These experiments were all conducted with the human K562 cell line.

We first set out to determine if RNase A treatment might lead to certain portions of the genome being underrepresented or overrepresented in the input DNA in the rChIP protocol, which then might give artificial peaks or loss of peaks when plotting ChIP/Input. Input DNA is the most commonly used control in ChIP and controls for biases in variable solubility of chromatin, chromatin shearing and amplification (Park 2009). To test for changes in genome coverage in the input DNA in the rChIP-seq protocol, we treated the input chromatin with RNase A and compared it to chromatin that was untreated, both in biological triplicates. Coverage in the six input samples appeared to be similar (Fig. 2A). We then used MACS2 software (Gaspar 2018) to identify regions of DNA that are different in RNase treated versus untreated input chromatin. MACS2 called only 6 significant regions of difference. This is in contrast to the hundreds of differentially represented sites in PRC2 rChIP experiments. Thus, the global representation of input chromatin is similar in RNase-treated and non-treated samples and does not reveal differential loss of chromatin upon RNase A treatment.

Next, we wanted to determine if the rChIP results with a particular protein-antibody combination would be reproduced in a different cellular setting. We had previously determined that ChIP of RBP1, the catalytic subunit of RNA polymerase II, was not substantially affected by RNase A treatment using our established rChIP protocol in IPSCs. Thus, we performed rChIP for RBP1 in K562 cells (Fig. 2B). We observed strong coherence between RNase-treated and untreated ChIP samples with very few regions showing differential representation (Fig. 2B,C). This result is expected for a protein that should not require RNA for its chromatin binding. Thus, the rChIP protocol applied to RBP1 gives concordant results in iPSCs and K562 cells, with no evidence for artifacts caused by RNase A treatment.

There are many chromatin complexes that are proposed to bind to RNA (Khalil, Guttman et al. 2009, Hendrickson, Kelley et al. 2016), and rChIP should be a useful approach to distinguish if RNA influences their localization or plays a different role. One such protein that specifically binds DNA and RNA is the CCCTC-binding factor (CTCF) (Saldana-Meyer, Gonzalez-Buendia et al. 2014, Hansen, Hsieh et al. 2019). Thus, we performed rChIP on CTCF using the same protocol as described in (Long, Hwang et al. 2020), using 5 replicates of CTCF ChIP and rChIP. We determined regions of significant CTCF binding, genome-wide, relative to three independent input samples using MACS2. Next, we used DeSeq2 to determine any peaks that were significantly changed in read density between RNase- and non-treated samples. We observed that CTCF bound to the same regions of DNA in the presence and absence of RNase (Fig. 2D, data shown in green). In fact, we only observed one DNA binding site that lost binding upon RNase treatment out of over 25,000 peaks identified in ChIP and rChIP conditions.

Next, we compared how RNase treatment would affect CTCF localization when applied at the IP stage, as in (Long, Hwang et al. 2020), versus the wash stage where only the immunoprecipitated protein, DNA and isolation beads remain; the latter modification should avoid potential precipitation issues caused by RNase treatment in the IP stage. To determine the global variability of CTCF binding sites identified in rChIP and ChIP we plotted the log2 fold DNA enrichment in RNase versus untreated conditions (Fig. 2D). The DNA enrichment followed a significant linear trend when RNase was applied at the IP (R = 0.94, Pval < 2e-16) or the wash stage (R = 0.93, Pvalue < 2e-16). Overall, this suggests that CTCF binds to the same DNA sites with the same representation in RNase-treated and untreated samples, independent of whether RNase is added in the IP or the wash stage (see also Fig. 2E).

### Improvements in the rChIP protocol

One disconcerting feature of our original rChIP-seq protocol was that the amount of DNA pulled down with the magnetic antibody beads was considerably higher in the samples subjected to RNase A treatment, as also reported by (Healy, Zhang et al. 2023). In the example shown in Fig. 3A, which concerns EZH2 ChIP with iPSC lysate, the DNA pull-down with RNase A was about 6-times higher, but this was variable between different antibodies and between different days of experimentation. It seemed that RNase A, being a positively charged nucleic acid-binding protein, might be binding to DNA and causing it to pull down with the beads. We therefore tested lower RNase A concentrations in the rChIP method detailed by (Long, Hwang et al. 2020). The lower concentrations of RNase A still gave complete digestion of total iPSC RNA (Fig. 3B), and they largely eliminated the pulldown of excess DNA (Fig. 3A). Importantly, there were still ∼500 genes whose pulldown by the anti-EZH2 antibody was significantly reduced by this low RNase A treatment (Fig. 3C), consistent with RNA bridges or tethers between PRC2 and chromatin. Quantitatively, the reduction in pulldown by low RNase A was smaller than the almost complete abolition of pulldown reported for high RNase A (Long, Hwang et al. 2020).

**Figure 3.**
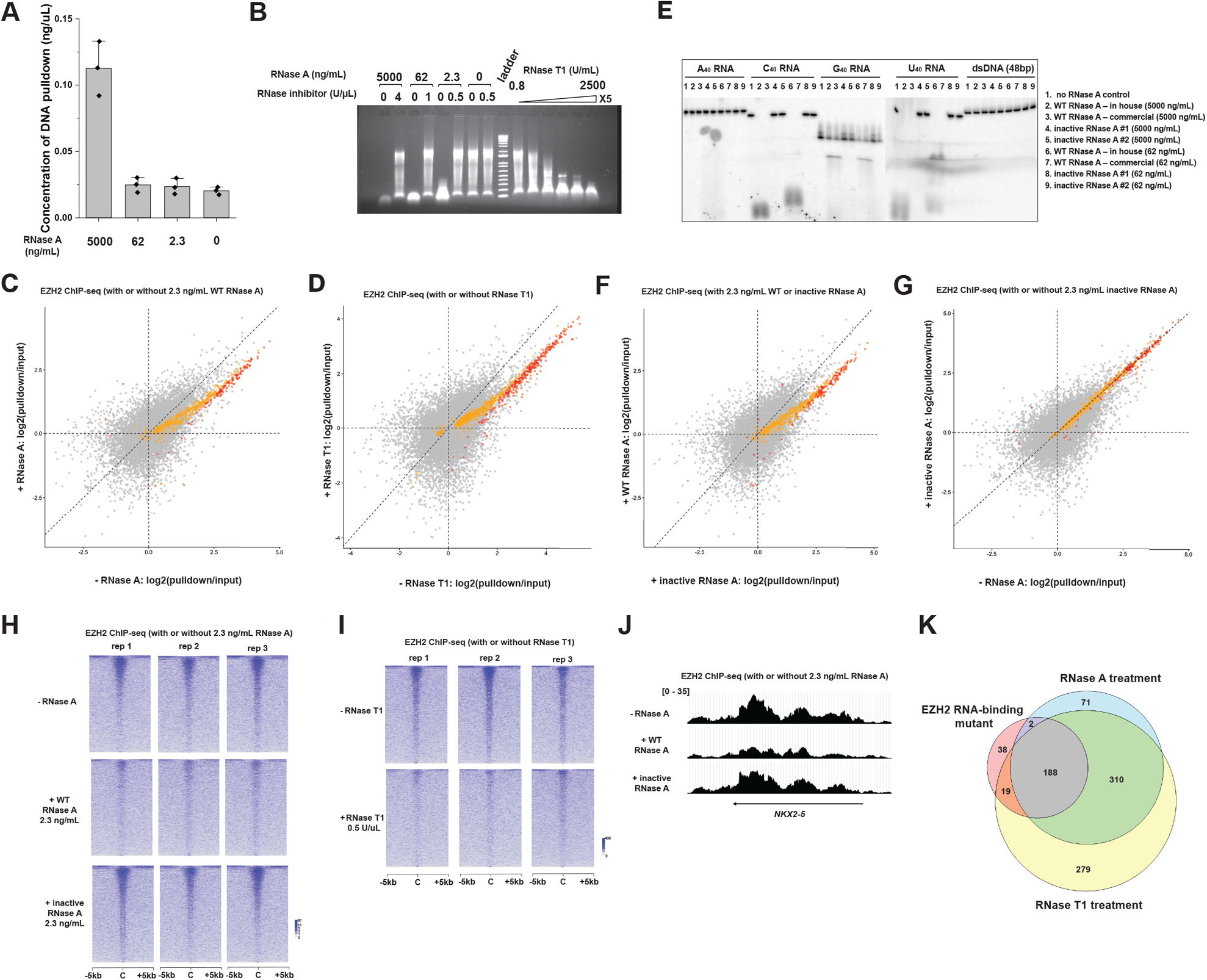
rChIP-seq optimization. **(A)** DNA yield after EZH2 rChIP-seq experiments using different concentrations of RNase A: 5000, 62, 2.3 or 0 ng/mL. 5000 ng/mL is the original concentration used in Fig.1 of (Long, Hwang et al. 2020). All ChIP pulldown DNA was resuspended in 40 μL of TE buffer. Three biological replicates were performed using crosslinked lysates of human induced pluripotent stem cell (hiPSC) line WTC-11. Error bar represents the standard deviation. **(B)** Effectiveness of RNase A and RNase T1 in digesting total RNA from hiPSC line WTC-11. 2 μg of total RNA from hiPSC was treated with different amount of RNase A with or without the RNase inhibitor (left side of the ladder), or a serial titration of RNase T1 from 0.8 Units/mL to 2500 Units/mL (5-fold increase, right side of ladder). RNase treatment overnight at 4°C in the ChIP IP buffer to approximate the rChIP-seq conditions. The digested RNA samples were loaded onto a 1% agarose 1XTBE native gel with ethidium bromide. 2.3 ng/mL of RNase A and 500 U/mL of RNase T1 was sufficient to completely digest almost all RNA, and these concentrations were used for downstream rChIP-seq experiments. **(C-D)** Gene scatter plots of EZH2 rChIP-seq experiments using 2.3 ng/mL RNase A and 500 U/mL RNase T1. Gray dots are insignificant comparing the X and Y values, yellow dots are significant (C: FDR<0.05, D: FDR<0.10) and log2Foldchange Y/X < 1, and red dots are significant and log2Foldchange Y/X > 1. **(E)** Wild type (WT) and catalytically inactive mutant of RNase A were expressed in *E*.*coli*, purified and tested for cleavage of 5’-^32^P radiolabeled A_40_, C_40_, G_40_, U_40_ RNA or a 48-bp double-stranded DNA under same conditions as in panel B. Treated nucleic acids were resolved on a 10% acrylamide, 7M Urea, 1XTBE denaturing gel. RNase A inactive mutant contains the triple mutation H12A/K41A/H119A. **(F-G)** Gene scatter plot of EZH2 rChIP-seq comparing with and without the inactive RNase A mutant. For panel F, Gray dots are insignificant comparing the X and Y values, yellow dots are significant (FDR<0.10) and log2Foldchange Y/X < 1, and red dots are significant and log2Foldchange Y/X > 1. For panel G, the grey, yellow and red coloring of each gene is based on panel C, to show that the mutant RNase A no longer causes loss of EZH2 on chromatin as shown in C. Note that these significantly changed genes in panel C are all lined up in the Y=X diagonal dash line in panel G. **(H)** Heat map of called peaks in EZH2 rChIP treated with 2.3 ng/mL RNase A (WT or inactive). Peaks are centered with the flanking 5kb genomic region and sorted by intensities. **(I)** Heat map of the called peaks in EZH2 rChIP treated with RNase T1. Generated in the same way as panel H. **(J)** A representative genome trace of EZH2 ChIP-seq showing the *NKX2-5* gene locus. **(K)** Venn diagrams with number of differentially occupied genes in EZH2 ChIP-seq experiments. Pink circle: differentially occupied genes between the WT and RNA-binding mutant EZH2; blue circle: differentially occupied genes between 2.3 ng/mL RNase A treatment and no treatment; yellow circle: differentially occupied genes between 500 U/mL RNase T1 treatment and no treatment.

We next tested RNase T1, which cleaves RNA after G residues instead of the pyrimidines. RNase T1 also degraded iPSC RNA to small fragments (Fig. 3B) and reduced the pulldown of ∼800 genes, most of which are PRC2 target genes (Fig. 3D).

One potential shortcoming of the rChIP-seq protocol is that experimental samples are being treated with a basic nucleic acid-binding protein whereas control samples are mock treated. It seemed that adding a catalytically inactive mutant of RNase to the control samples might give a more appropriate comparison. We first expressed both the wild-type RNase A and the inactive mutant (H12A/K41A/H119A, amino acid numbering based on the human homolog RNase 1 (Ressler, Mix et al. 2019)) in E. coli, purified the proteins, and assayed their activities. The WT RNase A cleaved poly(C) and poly(U) but not poly(A) or poly (G), as expected from its known specificity (Fig. 3E). In contrast, the inactive RNase A did not cleave any of the substrate RNAs (Fig. 3E). In genome-wide EZH2 ChIP-seq, using the inactive RNase A as the control sample gave the same results as when mock treatment was used as the control (compare Fig. 3F with 3C). Direct comparison of inactive RNase with mock treatment showed that they were equivalent (Fig. 3G). Thus, the conclusion that PRC2 occupancy of a subset of genes is mediated by RNA still holds if catalytically inactive RNase is used as the control.

Heat maps of called peaks showed a diminution of signal when rChIP-seq was performed with the low RNase A concentration but not when inactive RNase A was substituted (Fig. 3H). Heat maps also showed a diminution of signal when rChIP-seq was performed with RNase T1 substituted for RNase A (Fig. 3I). A representative genome browser trace for *NKX2-5*, a gene for a transcription factor involved in heart development, shows the typical broad distribution of PRC2 across the gene with peaks diminished upon low RNase A treatment but not by inactive RNase A (Fig. 3J). Finally, the genes with EZH2 occupancy significantly reduced by low RNase A treatment overlapped substantially with those reduced by RNase T1 treatment or by the RNA-binding mutant (Fig. 3K). Thus, our new analysis recapitulates our previous conclusion that RNA contributes to PRC2 occupancy on chromatin, although the effect size is more modest with the reduced concentration of RNase A or with the substitution of RNase T1.

Our conclusions about RNA contributing to PRC2-chromatin binding are also consistent with studies using the transcription inhibitors triptolide (Extended Data Fig. 2 of (Long, Hwang et al. 2020)) and DRB (Wei, Xiao et al. 2016). In both studies, treatment of transcription inhibitors led to loss of PRC2 on PRC2-target genes and gain of PRC2 on non-target genes. These drug protocols do not involve RNase A, so the similarity of these data with the rChIP-seq data argues against a substantial RNase A artifact in rChIP-seq.

In conclusion, our optimized rChIP-seq approach confirms our original finding that RNA contributes to PRC2’s chromatin occupancy, although the magnitude of the RNA contribution appears smaller in the new data presented here. For other researchers interested in rChIP-seq, we recommend using a low concentration (e.g., 2.3 ng/mL) of RNase A, preferably comparing the wild-type RNase and the catalytically inactive mutant. Treatment with another RNase such as RNase T1 is also recommended. Finally, transcription-inhibiting drugs can be used to interrogate the dependence of protein-chromatin binding on nascent RNA as an orthogonal method that uses no RNase.

## Acknowledgements

We thank Chen Davidovich (Monash University, Melbourne, Australia) and Richard Jenner (University College London, London, UK) for sharing their concerns about our prior publication. We thank Ron Raines (M.I.T.) for discussion of mutations that inactivate RNase A and for RNase A expression plasmids. Y.L. was supported by NIH K99 award no. K99GM132546. T.H. was supported by NIH K99 award no. K99GM137072. J.L.R. was a Howard Hughes Medical Institute (HHMI) Faculty Scholar. J.L.R. and T.H. were supported by NIH P01 award no. P01GM099117. T.R.C. is an investigator of the Howard Hughes Medical Institute (HHMI).

## Data availability

Genomic data is available at GEO accession GSE128135

